# What Helps Interface Projects in Ecology Work? Learnings on Facilitating Conditions from a Decade of Inter- and Transdisciplinary Research on Invasive Fish

**DOI:** 10.1101/2025.05.05.652198

**Authors:** Irene Adrian-Kalchhauser, Karen Bussmann-Charran, Anouk N’Guyen, Philipp Emanuel Hirsch, Joschka Wiegleb, Lukas Bergmann, Patricia Burkhardt-Holm

## Abstract

Ecological research for environmental management occupies a unique and demanding space at the interface of societal needs and academic research. Projects in this space must reconcile stakeholder involvement and real-world applicability with academic requirements such as disciplinary excellence and career advancement.

Here, we present a post-hoc account of a 12-year inter- and transdisciplinary research initiative on the management of invasive gobies in central Europe. Based on a synthesis of inputs, outputs, and outcomes, we distill facilitating conditions that increased the likelihood of the project reaching its goals across political, administrative, societal, institutional, team, and individual levels. Rather than “success factors” in a deterministic sense, these are conditions and practices that created a favorable environment for both academic and applied outcomes. The interaction of these domains created windows of opportunity that could be seized for both scientific progress and societal impact.

By reflecting from the perspective of natural scientists directly engaged in such a project, we aim to complement existing social science frameworks with an insider view of the lived realities of inter- and transdisciplinary collaboration. Our retrospective can inform policy makers, funders, and institutions seeking to create enabling environments for interface research, and may support fellow natural scientists in proactively preparing the ground for their own initiatives.

## Introduction

### Among the natural sciences, research activities gathering data for environmental management strategies occupy a unique and challenging realm

The expectations of researchers, funders, and stakeholders with regards to research objectives, scientific rigor, and communication are often distinct (Lawrence et al. 2022). Society and policy call for swift progress and unequivocal, locally obtained results, which misaligns with the measured pace and circular process of scientific investigations, their goal of producing deliverables supporting scientific careers, the expectations of monodisciplinary “excellence” of funding agencies, and the inherent intention of science to advance scientific knowledge beyond individual or local case studies. The production of natural sciences knowledge that is applicable to environmental management while meeting the requirements of fundamental academic research is thus non-trivial and requires additional efforts (Deutsch 2025).

### The field of invasion biology is particularly rich in projects designed to both inform environmental management and uncover fundamental biological truths

Within this domain, the co-production of knowledge and the integration of natural and social science disciplines represent the state of the art (Bessert-Nettelbeck 2023). Because invasive species serve as nucleation points for novel interactions between nature and society, invasion biology is a domain in which socio-ecological knowledge generation systems are particularly frequent (Kemp et al. 2017). Projects in this realm frequently qualify as “transdisciplinary”, since they rely on the collaboration of researchers and practitioners from the private and public sectors of civil society in problem definition and knowledge production towards societal problem solving (Pohl 2021). Procedural insights obtained from invasion biology projects may therefore offer valuable lessons, for example for funding agencies that are active in this area, and may provide conservation managers with a deeper understanding of the temporal and group dynamics and requirements of applied research projects (Deutsch 2025). Invasion biology projects may also serve as compelling examples on how to stimulate discussions and foster comprehension among the environmental managers, funders, and researchers of tomorrow (Pohl 2021).

### Various factors that create a favourable environment for such projects, as well as measures of success, have been defined previously by the social sciences

Conducive conditions include in-person engagement and a desire to truly learn on the individual level, an awareness of “hidden” agendas at the team level, the budgeting of time for integration on the program level, the recognition of the value of applied research at the institutional level, and the willingness to navigate around technocratic hurdles on the socio-political level (Deutsch 2025). Success factors useful for formative evaluations of transdisciplinary research have been derived from the study of real-world laboratories (Wiefek 2024). These are inherently multi-dimensional along the spatio-temporal scales and various actors’ perspectives, and can include natural science research outputs, such as peer-reviewed publications or graduations, tangible outcomes such as the development of new management strategies, or the implementation of relevant policies and legal frameworks. Another dimension of evaluation is the collaborative process itself and the effective integration across disciplinary and societal boundaries (Pohl 2021). The quality of stakeholder involvement, the creation of mutual trust, and the establishment of sustainable networks and new forms of cooperation are difficult to quantify, yet crucial for productive cross-boundary knowledge production. Finally, the ability to foster learning among scientists, practitioners, and stakeholders alike is a component promoting long-term impact (Pohl 2021), since success in invasive species research is not so much about actually solving a problem (measures are usually implemented, or not, by non-academic actors, with policy decisions taken at higher levels), but more about building capacity across sectors. For transdisciplinary projects, success has thus been defined as a) improvement in the situation for both researchers and practitioners, b) production and dissemination of artefacts that contribute to stocks and flows of knowledge, and c) mutual and transforming learning for both researchers and practitioners (Pohl 2021).

### Here, we present an example case of an interface project

We do not aim to engage in a formative evaluation with regard to sustainable solutions to societal problems sensu Wiefek (2024). Rather, we intend to use the exemplary case of a problem-driven, multi-year, inter- and transdisciplinary research initiative to share the perspective of the natural science research partners involved. While the literature is replete with social science accounts studying inter- and transdisciplinary projects, the involved natural science researchers rarely take a meta-perspective themselves. Our expertise on the overarching socio-ecological systems is explicitly limited, but we believe a peer-to-peer account aimed at natural scientists stumbling into these systems through their research interests will add a valuable perspective, particularly for natural science research teams. In addition, the account can be used as a study and teaching example for lecturers and students in the field of environmental management.

### The project under consideration unfolded in Switzerland from 2012 to 2022, and focused on invasive fish

“Non-native goby species in Switzerland – Measures to control and minimize their impact” was sparked by the detection of invasive gobies in the Rhine in Basel in 2011 during routine governmental monitoring of juvenile fish along the riverbanks. Consequently, local researchers, regional stakeholders and federal administration launched an initiative to define problem areas, elucidate the ecological context of the invasion and to propose and test measures to stop further dispersal. Specific objectives were defined collaboratively, and included finding the source (identify origin and introduction pathways), monitoring the population (follow local population dynamics and the expansion of geographical distribution), understanding dispersal (uncover natural and anthropogenic factors facilitating dispersal beyond the introduction site), assessing impact (evaluate the impact of the invasion on the native fauna), devising prevention strategies (suggest effective measures to prevent dispersal and mitigate impact), and communicating and educating (inform, raise awareness, and educate relevant target groups about the risk of human-assisted dispersal). Importantly, the *implementation* of measures was not part of the research project; the aim was to *inform* relevant stakeholders about their options. Nonetheless, the project could be considered inter- and transdisciplinary sensu Pohl (2021), specifically a “problem-solving setting with methodological, mostly “intermediate-scope” interdisciplinarity, with a-priori awareness for the complexity of the issue and the diversity of perceptions, aiming to link abstract with case-specific knowledge to propose solutions for a more sustainable development.” Practice and science came together from the outset for problem framing, problem analysis, and impact explorations.

The account encompasses a description of the context, of the input in terms of human and financial resources (**Fig. 1**), a comprehensive overview of the outcomes across three sectors (**Fig. 2**), a description of the contribution of stakeholders to key results (**Fig. 3**) and most crucially, a set of internal and external conditions - from political system to personal values of team members - that in some way impacted the project positively. These were retrospectively and subjectively distilled from the project by the authors through discussions and iterative revisions of a draft document (**Table 1**).

**Table 1.** Factors facilitating and supporting id/td research projects.

| Project-external factors (settings the project takes place in and the research team has no control over) |  |  |
| --- | --- | --- |
| Level | Factor & Description | Examples & Situation in the Goby Project |
| Political level | **Conducive political culture.** The local and federal political environment grounds decisions in evidence and is willing to translate research results and suggestions of experts into practice. | Swiss political culture overall values evidence and experts’ inputs. Local political representatives are in close touch with public opinion and with key stakeholders. On some occasions, this facilitated action: a prototype in-stream barrier was indeed installed. On other occasions, conflicts of interest were stronger than evidence: measures impacting individual freedom (e.g. boat cleaning) were not widely implemented. |
|  | **Topicality and social concord:** Democratic processes and legislative initiatives in the subject area have previously set societal goals and boundaries, clarified shared values, and highlighted knowledge gaps that guide future research needs. | Coinciding with the project, the National Biodiversity Strategy (2012) and the updated Fisheries Ordinance (2021) mandated action against invasive species, while the Federal Water Protection Law and Ordinance required the renovation of fish passes to ensure fish migration (SRF 2024a). Swiss politics and administration therefore had prior knowledge regarding invasive species and associated ethical and political challenges, and there was minimal dissent on the issue’s importance. |
| Administrative level | **Approachability.** Federal and cantonal administrations are approachable, personally reachable, and actively support the bidirectional flow of information. | Administration representatives were available for personal discussions, actively invited inputs (e.g. invitations to present the state of the art in national invasive species working groups), and communicated new regulations or changes in processes. Research plans and proposals submitted to funding could be candidly discussed with administration prior to submission. Obtaining permits for sampling was straightforward. |
|  | **Decentralized Policy Responsibility:** Local (approachable) administrative units hold decision-making power over funding, collaboration, and project support. Time to decision is therefore short, and collaboration is efficient. | In federalist Switzerland, local authorities hold significant decision-making power as well as leeway in harm-benefit analyses and can therefore come up with workable solutions quickly. For example, the protected nase must not be disturbed during spawning season. To maintain legal requirements while enabling crucial sampling, the local fishery inspectorate was personally involved in designing and executing the field sampling to ensure conservation goals and research needs were both met. |
|  | **Diverse Funding Landscape:** Diverse funding opportunities are available across the axes of applied to basic research, of short-term activities to multi-year stipends, and of private to | The 10-year course of the project was supported by a colorful diversity of grants supporting both fundamental and applied research. For instance, the Swiss National Science Foundation funded work on characterizing aspects of the round goby genome, while a consortium of cantonal |
|  | public funders. | funds supported an applied study on the role of recreational boating in round goby translocation. |
| **Societal level** | **Consensus culture:** The societal and political culture is used to dialogue and compromise, attempts to find consensus when facing emerging challenges, and listens to scientific evidence. | In Switzerland, a deeply rooted culture of dialogue, compromise, and consensus seeking yields broadly accepted solutions, even when policies have conflicting goals. For example, the water protection law requires all waterways to be passable for all fish species by 2030, while the national strategy on invasive alien species calls for barriers to the spread of invasive species, which led to solution-oriented discussions between policymakers, administrators, and scientists to strike an acceptable balance. While a legal mandate to clean boats did not find support (due to a very high regard for personal autonomy), in a practical compromise, scientists, water sports clubs, and authorities collaborated to install a voluntary cleaning station. |
|  | **Value of study object or associated resources:** Research projects targeting a subject that has high value to society (e.g. economical, recreational, symbolic) benefit from public attention and can create interest and discourse for the problems and solutions at hand. | Switzerland harbors the springs of four major European watersheds. The protection of water bodies is a national priority, both for their ecological importance and symbolic value. Significant resources are invested in aquatic conservation, such as salmon restocking programs. Public interest in water and nature-related issues is strong, which in our case lead to invited talks at stakeholder organizations, media coverage in newspapers and topical journals, and facilitated project funding. |
|  | **Established topical networks:** Pre-existing stakeholder networks offer opportunities to connect science with society, and advance research by providing access to locations, resources, local knowledge, and feedback on proposed measures. However, tight-knit club-like networks can also be challenging to access for outsiders. | Switzerland features a strong culture of civic participation through associations, clubs, and collaborative structures. This led to institutionalized interest in the research project and facilitated the integration of members of the public in transdisciplinary processes (N’Guyen et al., 2016, Hirsch et al., 2020). At the same time, this created a bias towards individuals organized in such associations. Also, reaching groups who are rarely organized in formal associations, e.g. aquarists, proved challenging, and entering networks dominated by senior males was at times daunting for junior female team members. |
| **Institutional level** | **Institutional commitment and structures:** id/td projects benefit from academic evaluation systems that recognize societal impact and offer access to dedicated structures such as knowledge brokers and digital platforms. Other beneficial institutional aspects are training opportunities in communication, facilitation, and stakeholder engagement, and flexible administrative and legal frameworks that support unconventional project setups. | The institute hosting the project is also hosting an interdisciplinary master’s program (Master of Sustainable Development). This facilitated the contact between natural and social scientists as well as economists and the recruitment of students from different backgrounds with an interest in projects that cross disciplinary boundaries. It also reliably receives a stable amount of personnel funds, disposable funds, and equipment funds from the department it is associated with, which facilitates initial explorations prior to funding acquisition. |
| **Other** | **The Perfect Storm of Opportunity:** The convergence of favourable conditions can create powerful momentum, and the combined effect of several conducive elements then exceeds what any single factor could achieve alone. | Two such dynamically interacting factors in the project were the geographical proximity between research objects, stakeholders and researchers, the strong identification with affected water bodies, and the presence of exceptionally engaged individuals in decision-making capacities. |
| **Project-internal factors (settings and decisions the research team has control over)** |  |  |

| Level | Factor & Description | Examples & Situation in the Goby Project |
| --- | --- | --- |
| **Political level** | **Awareness of Political Context:** Actively aligning research plans and activities with national and international strategies, policies, and regulations, and an awareness of shifting priorities, increases the likelihood that research findings will be considered by policymakers and practitioners, and translated into science-based action. | The project was designed with careful consideration of key regulations related to invasive species and water management in Switzerland (see above), ensuring that research strategies and experimental approaches were aligned with practical policy needs. From the outset, the team maintained a strong awareness of ongoing policy developments, and all members were expected to stay informed and responsive to the evolving regulatory context. |
|  | **Grooming the science-policy interface:** A consistent engagement at the science-policy interface and proactive management of good connections with governmental and administrative bodies fosters mutual trust and saves time during planning and implementation of new projects. | Previous activities of the project lead at the science-policy interface allowed the project to build on existing relationships and zones of contact between science and politics. Consequently, the project started from a high level of trust and recognition by authorities, as exemplified by multiple invitations to consult on policy matters and contributions to submission of parliamentary interventions, which in turn became important drivers of the project's visibility and influence. |
| **Administrative Level** | **Process awareness:** All project partners must be able and willing to engage in open and upfront communication about their own explicit and implicit expectations and limitations of scientific methods. For example, policymakers tend to seek quick, definitive high certainty answers that science cannot reliably provide. | Limited resources and data made a full scientific risk assessment unfeasible. The team thus developed recommendations for preventive measures coupled with scenarios related to non-implementation. During stakeholder events, participants then used Likert scales to assess the likelihood of success of proposed measures and thus prioritized actions despite uncertainties. |
| **Societal level** | **Cultural awareness:** Scientists are often immigrants to their country of residence and work. An awareness and open discussion of cultural norms, communication styles, and institutional practices improves science-society exchanges. | Cultural differences and the need for approaches that would not be everyone's primary personal choice were openly discussed and reflected. This included for example the introduction to the concept of the professional Apero as key ingredient of informal knowledge exchange and trust building. |
| **Institutional level** | **Institutional support:** units doing id/td research contribute to different reputation aspects than units performing monodisciplinary science. Continuous efforts to maintain institutional commitment are key for long-term interface projects. | The team took active steps to raise the visibility and to underline the relevance of their work within the framework of available vessels. This included active participation in department meetings, presentation of results at colloquia, engagement as postdoc representative, or participation in lecture series and in outreach. |
|  | **Teaching feedback:** Integrating id/td research projects into teaching enhances theoretical grounding through engagement with the literature, supports student recruitment, and benefits from fresh input and perspectives contributed by students. | The goby project was repeatedly used as an example in various courses about ethics, ecology, and aquatic biology. It presented a tangible case of high interest for students in the Master of Sustainable Development considering the relevance of invasive species across social, economic, and scientific dimensions. |
| **Project level** | **Shifting Priorities and Resource Flexibility:** All research projects start with a plan, but id/td projects are more likely than others to change course. The ability to accept shifted priorities and to adapt to changed resource availability is therefore key. This includes aspects of team composition | Project components related to ballast water and boat cleaning turned out smaller in scope than originally planned, despite very clear relevance, partly due to stakeholders' opinions and partly due to obstacles related to data collection and accessibility. Another potentially important line of inquiry related to anti-fouling agents was not pursued despite an initial planning and research stage due to a lack of resources. |
|  | (individual strengths, interests, and limitations) that may not align with initial goals, and require frequent strategic checkpoints. |  |
|  | **Proactive Networking:** Treating networking as an integral project component and a primary task fosters mutual understanding, builds trust, and helps align goals and expectations. By embedding networking into the project's structure from the beginning, researchers can better identify synergies, anticipate challenges, and respond flexibly to emerging opportunities. | The project emphasized proactive, structured networking with stakeholders from the outset. Research priorities were defined jointly to consider both scientific goals and practical needs. Tailored annual reports addressed stakeholders' evolving questions and priorities, annual exchange events further strengthened these relationships and offered space to share updates and actively solicit feedback and input. The resulting two-way dialogue enriched the research and its real-world application. |
|  | **Community Engagement Beyond Formal Roles:** Building genuine connections with stakeholder communities leads to deeper insights, unexpected collaborations, and builds trust. Taking the time to engage informally, often outside regular working hours can be resource-intensive and personally demanding but is quite essential in id/td research. | Spending time with local fishing and boating communities and seeking out informal, unplanned encounters led to several unexpected yet highly productive collaborations on boat motor use and ballast water systems, and yielded access to sampling sites, samples, and sighting reports. These commitments were not solicited by team members, but were always welcomed. Team members were always encouraged to seize interaction opportunities when they arose, with travel expenses reimbursed and time investments recognized. |
| **Team level** | **Active management of team diversity.** Diversity in disciplinary backgrounds, professional expertise, and personal perspectives is at the core of id/td research but must be actively managed. Beyond the conscious assembly of complementary skills and viewpoints, this includes deliberate efforts to create an environment where conflict is recognized, tackled, and managed so that differences can become a source of creativity rather than of misunderstanding and frustration. | Conflict was allowed, but simultaneously, team-building exercises cultivated mutual appreciation for the contrasting disciplinary backgrounds. Regular opportunities for meta-level reflection on team dynamics, e.g. distinct needs in terms of support and milestones (e.g. the differing scopes and timeframes for master and PhD theses) existed. Annual two-day retreats provided dedicated time slots to discuss contrasting perspectives and collaboration processes. |
|  | **Integrated career planning:** In id/td projects, it is essential to ensure that individual career development is not compromised by the overall project strategy. To address this, an open and ongoing dialogue within the team about career goals, publication strategies, and the specific demands of ID/TD research is crucial. | A practical approach adopted by the team was inclusive co-authorship. Team members were encouraged to mutually invite each others' contribution and to offer other team members co-authorship opportunities, provided their involvement met ethical authorship standards (intellectual contribution). Invitations were tailored to individual strengths and availability, such as preparing figures, conducting literature reviews, or drafting and integrating discussion sections. |
|  | **Willingness to collaborate:** A collaborative culture marked by structured resource sharing, mutual support, and the recognition of diverse perspectives as beneficial is essential. A collectively positive attitude toward internal collaboration and joint problem-solving promotes success. | Senior researchers actively onboarded new team members e.g. by establishing contacts with administrative units and stakeholder networks. The team invested in internal communication infrastructure, such as shared contact tables, an wiki for knowledge sharing, and an annual two-day retreat. Information relevant for other team members, e.g. from conferences, was actively collected and shared. |
| Individual level | **Curiosity, Openness, and Mental Flexibility:**<br>A mindset that embraces curiosity, intellectual freedom, and adaptability supports id/td work. The ability to maintain a “beginner’s mind”, for playful exploration, and the freedom of thought to treat science as both inquiry and creative process is essential. Mental flexibility, the ability to compromise, shift perspectives, and adapt plans is equally important. The ability to spot and grasp opportunities for one own’s as well as colleagues’ research interests also plays a role. | Team members were free to pursue unexpected or random questions and ideas. Time and resources were made available for exploratory efforts with a “try and fail” mindset. This approach led to DIY spawning trap construction, egg desiccation experiments, edibility tests, or hydraulic experiments with preserved goby. These activities generated surprising insights or, alternatively, led to rapid changes in direction, and helped build a culture of curiosity, experimentation, and innovation. This was selective on occasion and led to the loss of team members with different approaches to research. |
|  | **Transparency and awareness about career aspirations:** id/td work requires time for activities other than classical academic career-advancing measures and outputs. An awareness of own needs, expectations, id/td requirements, and a proactive navigation of the associated challenges is essential. | Regular open discussions around career development and the challenges of publishing interdisciplinary work were integrated into grant planning, project design, and publication meetings and raised both by junior researchers and the PI. This ongoing dialogue enabled the team to identify tailored solutions that balanced individual career goals with the collaborative demands of ID/TD research. |

**Fig. 1:**
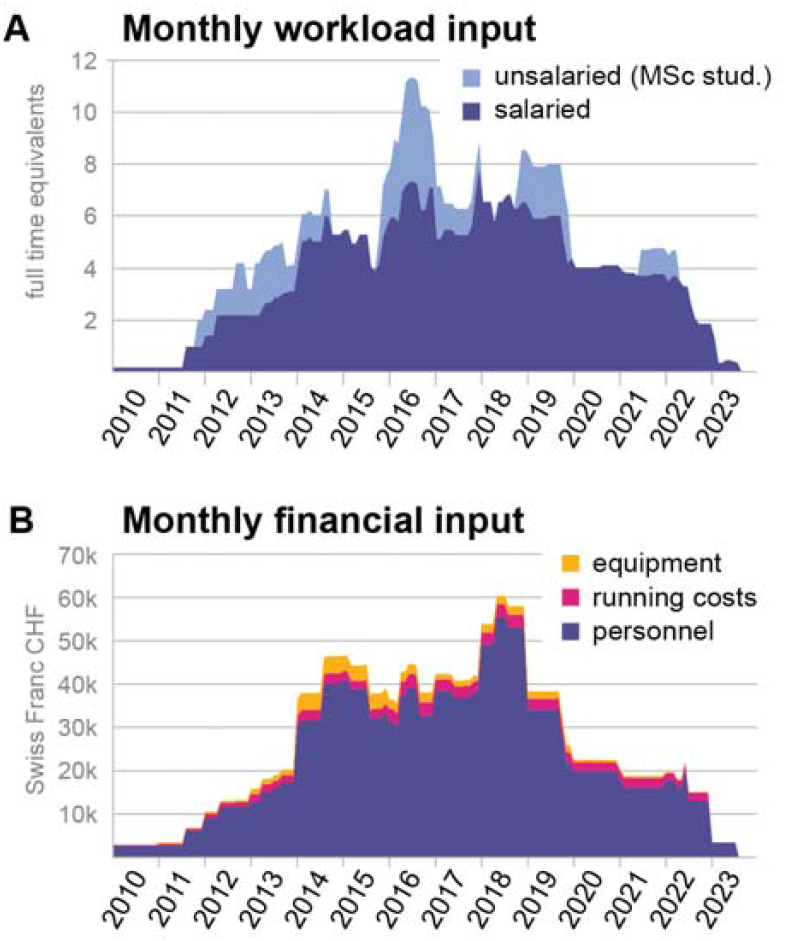
Resources consumed by the project. **A** Workload input per month follows a concave pattern over 12 years, with between 2 and 6 full time equivalents of paid personnel supported by 1-3 MSc students. **B** Financial input per month mostly follows the pattern of salaried personnel, with a non-scaling small fraction of running costs, and equipment investments peaking at 1/3rd of the project duration and trailing off in the last third of the project. During the core phase, the project required 40-50k in Swiss Francs per month, which mostly covered salaries. Non-salary components constituted 5% to 10% of total expenses.

**Fig. 2:**
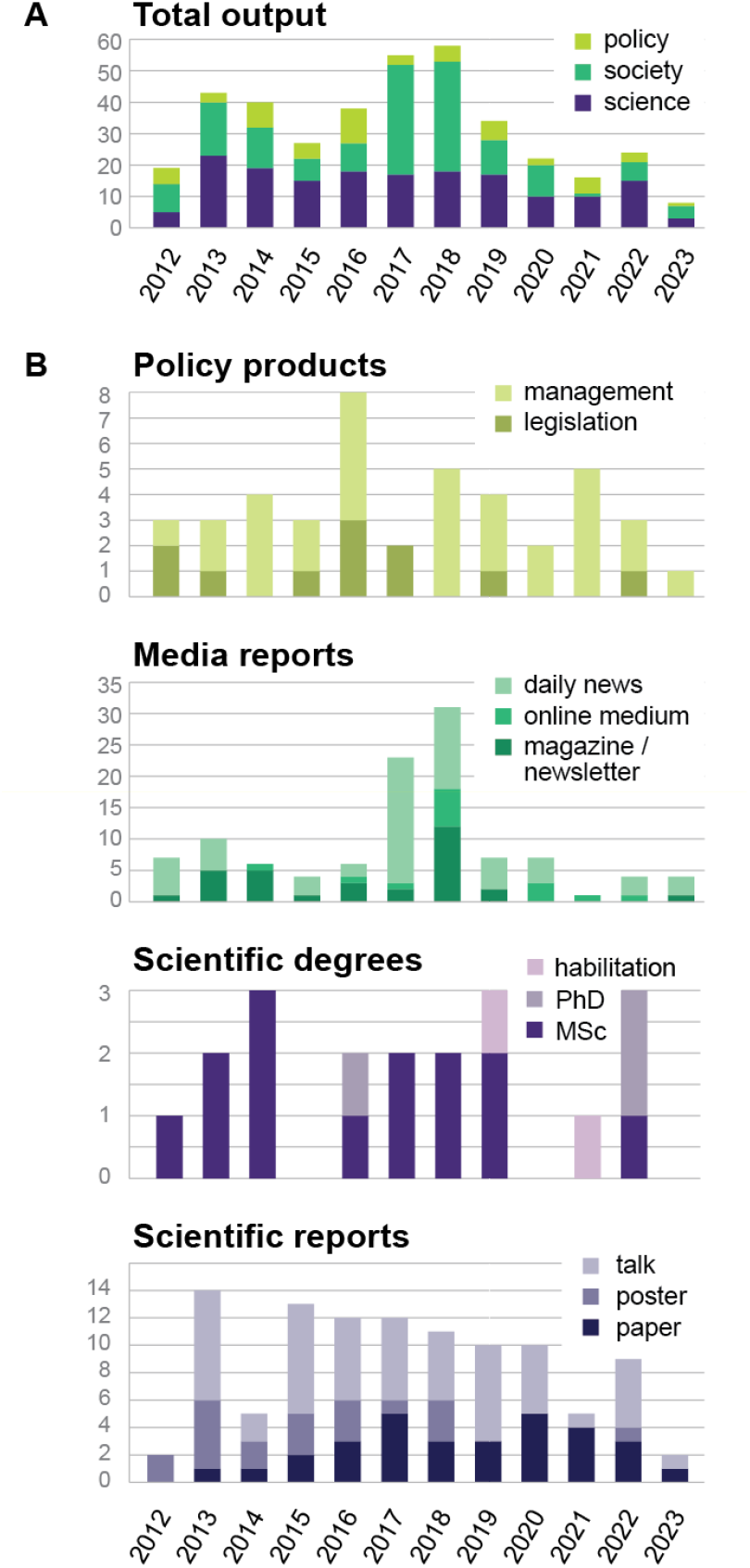
Project outputs. **A Total output** of the project across policy, society, and science (data from Supplemental Table 1). **B Policy products, Media reports, Scientific degrees, and Scientific reports**.

**Fig. 3:**
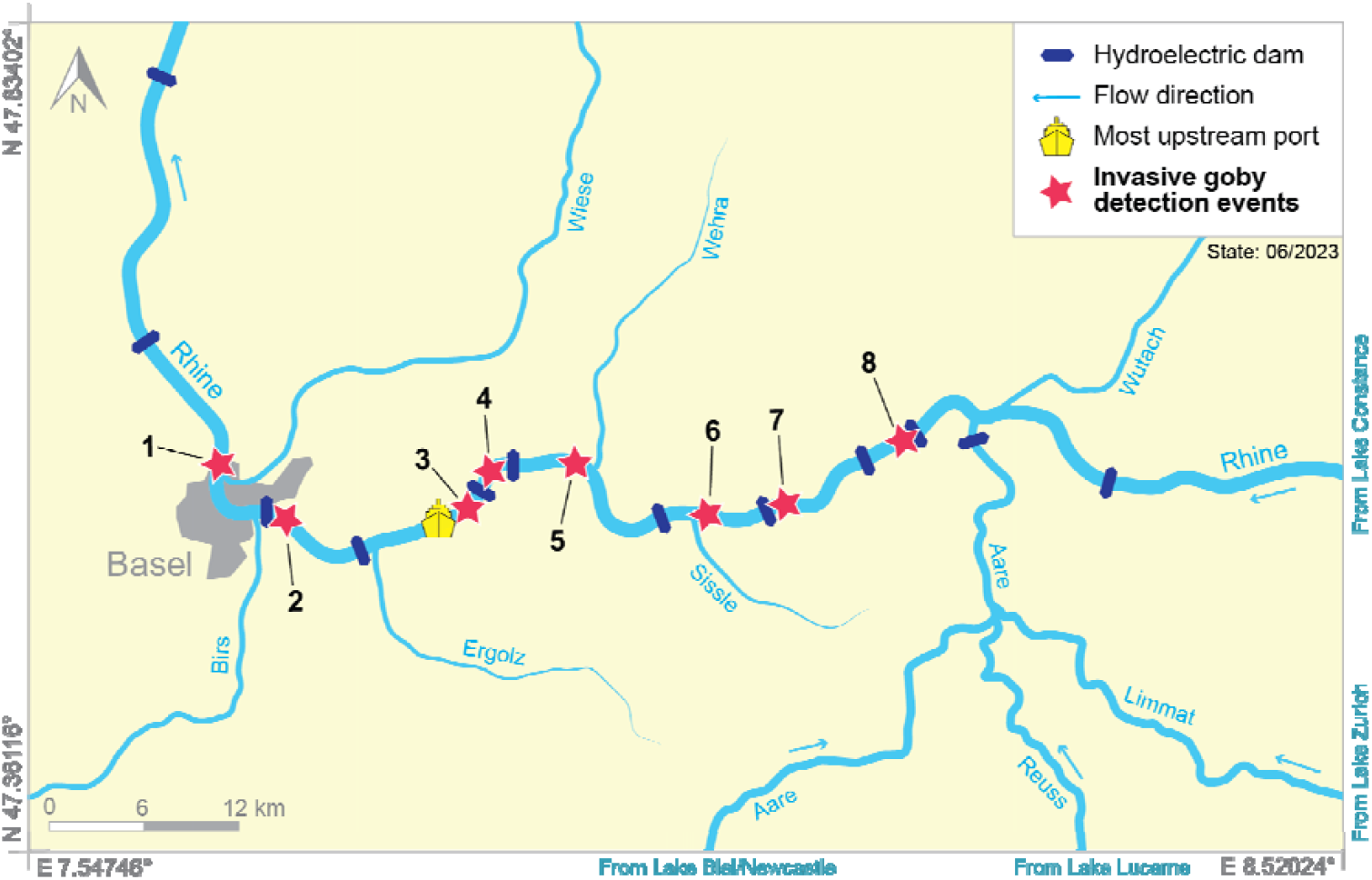
Dispersal of invasive gobies along the river Rhine. First detections upstream of the nearest hydroelectric dam (Numbered red stars) illustrate the role of the stakeholder network in monitoring the spread of invasive gobies in Switzerland. **1) Nov 2011** governmental electrofishing campaign. **May/Jun 2012** own minnow trap fishing campaign **2) Dec 2015** governmental monitoring campaign supported by local fishing communities. **3) Aug 2015** governmental monitoring campaign supported by local fishing communities. **4) Oct 2017** governmental campaign monitoring the efficiency of fish bypasses by ecobureau. **5) Aug 2017** governmental monitoring campaign supported by local fishing communities. **6) Sep/Nov 2018** snorkeler & local fisherman. **7) Jun 2020** local fishermen **Jun 2022** minnow trap campaign of cantonal authorities. **8) Jun 2023** local fishermen.

### Project characteristics and inputs

#### Team

Across 12 years, 16 academics were involved in the project, including 1 PI, 2 senior researchers, 5 postdoctoral fellows, 4 PhD students, and 4 master students, with an average of 3.7 full time equivalents across the entire project duration (**Fig. 1)**. 10 of these team members were female, and 7 male. The team additionally featured, in changing composition, 24 student assistants, 2 animal care takers, 5 administrative / IT assistants, and 2 lab technicians. First and last authors of 30 peer-reviewed papers had degrees in aquatic ecology, limnology, genetics, molecular biology, and sustainable development. Co-authors disproportionately included students with a master’s degree in sustainable development (an interdisciplinary masters offered by the host institution together with other faculties). External co-authors had affiliations in 8 different countries (Europe and US).

#### Policy and legislation landscape

Four notable policy works framed the project. In 1992, 20 years before the project started, Switzerland had signed the Convention on Biological Diversity (CBD) (United Nations 1992). In 2008, 4 years before the project started, the national fisheries ordinance was amended to mandate the prevention of the spread of and, if possible, eradication of alien fish and crustaceans, with federal administration in charge of coordination (SR 923.01, 2021) (Schweizerischer Bundesrat 24.11.1993). In 2012, when the project started, a national “biodiversity strategy” was adopted, stating the intention of Switzerland to conserve and promote biodiversity and ecosystem services for the economy and society (Bundesamt für Umwelt BAFU 2012). In 2016, four years into the project, a national “strategy on invasive alien species” further specified national regulations and international obligations regarding invasive alien species and identified necessary measures (Bundesamt für Umwelt BAFU 2016).

#### Funding portfolio

In total, the project required resources in the range of 4.4 Mio CHF^1^, of which 3.9 Mio CHF went into salaries, 0.3 Mio CHF into consumables, and 0.2 Mio CHF into infrastructure and devices (**Fig. 1B**). Funds were contributed by 5 different local administrations (0.36 Mio CHF), federal administration (0.4 Mio CHF), a collaborating university unit (0.2 Mio CHF), competitive research grants (0.33 Mio CHF), funding invested from a previous foundation contribution (0.39 Mio CHF), and the unit’s own annual funds (2.74 Mio CHF). In addition, the project used university infrastructure (offices, laboratories, seminar rooms, fish husbandry, storage) that are not monetized here.

#### Stakeholders and partners

Stakeholders invested in the project included practitioners such as anglers, water sports enthusiasts, boat owners, but also commercial shipping firms, port authorities, consulting firms, conservation organizations and authorities responsible for management of water courses and fisheries. The interaction relied on methods from the social sciences, such as Futures Wheels, questionnaires, in-presence workshops, expert interviews, transdisciplinary dialogues and scenario-driven discussions. Stakeholders contributed to data collection (for example, a network of cantonal authorities and fisher’s association was informally activated to report catch numbers of invasive gobies) and identified, prioritized and structured research targets (for example, the initial project prioritization of negative effects on native species and population control was co-developed with fisheries association and cantonal authorities; see N’Guyen et al. 2016).

#### Society & media

The project goals and outcomes were actively communicated by the involved researchers through various outlets **(Fig. 2C**). Targeted outreach activities such as annual on-site stakeholder workshops allowed for personal contact with invited media representatives, who were then proactively approached with new results and invited to join field work. Further outreach activities included presentations at local fishery societies, participation in outreach for schools and the public, visits to association headquarters, contribution to a famous Swiss children’s book series (Bieri 2018)) or the attendance of trade fairs.

### Project outcomes and outputs

The academic output of the project includes 29 peer-reviewed papers, 53 conference talks, 20 posters presented at conferences, 2 habilitations, 14 master theses, and 3 PhD theses (Fig. 2) along 5 major research axes: method development, spread, impact, adaptation, and microplastics. The project served as a stepping stone for a multitude of careers: the experience and scientific credentials gained allowed project team members to move into positions related to academic research and academic teaching, to teaching and communication positions, to applied research positions, and to positions dealing with nature-people interactions.

In accordance with the needs co-defined with various stakeholder groups, three axes of applied research were followed over the course of the project. First, novel methods were developed to meet the need for rapid detection of both the spread and the impact of the invasion. Detection methods evolved from standard bait-and-trap approaches (Kalchhauser et al. 2013) to the use of spawning traps (Adrian-Kalchhauser et al. 2018; N’Guyen et al. 2018), to eDNA based assays fine-tuned to the species and habitat characteristics (Adrian-Kalchhauser und Burkhardt-Holm 2016), and ultimately, methods to detect particular predation patterns of the species (Lutz et al. 2020). Despite this wealth of detection methods, however, non-academic stakeholders such as anglers ended up providing the most up-to-date information on the upstream spread of the fish (**Fig. 3**) - an indication for the tremendous value of integration.

Second, impact assessments and population models were developed to enable policy makers to take evidence-based decisions on management needs and -options while accounting for available resources. Literature review efforts found that the impacts of round goby were highly dependent on the ecosystem, time scale and life stage (Hirsch et al. 2016), and thus difficult to predict. In parallel, multi-year field-data were used to parametrize modeling approaches, which underscored the immense resource requirements for post-introduction management (N’Guyen et al. 2018). A key aspect of this research axis was the repeated discussion of concepts of risk management, event probabilities, scenarios and alternatives, and the value of precautionary management with stakeholders and administration.

Third, dispersal-related aspects were investigated to understand opportunities for preventative measures. These included active swimming, passive dispersal through boats and ships (Bussmann et al. 2021), active and passive dispersal through recreational activities, and active or passive dispersal through birds (not substantiated by the literature (Hirsch et al. 2018). Freshwater cargo ships and tankers were shown to be plausible long- and intermediate distance vectors based on population genetics (Adrian-Kalchhauser et al. 2016), egg desiccation tolerance (Hirsch et al. 2016), and use of harbour infrastructure (Bussmann und Burkhardt-Holm 2020; Bussmann et al. 2022; Adrian-Kalchhauser et al. 2017). Aspects such as personality traits (Hirsch et al. 2017) were also investigated, as was the ability of round goby to surpass in-stream barriers (Wiegleb et al. 2020; Egger et al. 2021), and the propensity for assisted dispersal by hobbyists and recreational activities (Hirsch et al. 2021). Results were immediately passed on to management and administration.

Observations made and samples collected for these three application-related research axes inspired additional investigations on adaptation and behaviour that went beyond the management-related research goals identified together with the stakeholders. These included the characterization of the round goby genome and mitogenome, the parental contribution in relation to temperature, and DNA methylation in relation to reproductive morphotype (Gutnik et al. 2019; Kalchhauser et al. 2013; Kalchhauser et al. 2016; Adrian-Kalchhauser und Burkhardt-Holm 2016; Adrian-Kalchhauser et al. 2017; Adrian-Kalchhauser et al. 2018; Adrian-Kalchhauser et al. 2020; Somerville et al. 2019), the capacity of the species for upstream dispersal inspired fundamental research into hydraulic properties and the physics of swimming behaviors (Egger et al. 2021; Wiegleb et al. 2022, 2023a; Wiegleb et al. 2023b; Govindasamy et al. 2024), and the species’ feeding behaviors and its ubiquitous occurrence inspired research into microplastic food chains and the potential of invasive gobies as indicators of benthic microplastic pollution (Bosshart et al. 2020).

The application-relevant findings were integrated into 31 non-academic management-related products, from leaflets for citizens to reports for administrative units, and 11 legislation products on the local, federal or international levels, such as parliamentary initiatives (**Fig. 2, SI**). On the political level, three interpellations, two postulates and one motion (Swiss parliamentary instruments) on the protection of aquatic biodiversity and the prevention of round goby spread referred to the project. The team’s activities resulted in the adaptation of the Ordinance to the Federal Act on Fisheries, namely, the inclusion of five species of Black Sea gobies in the list of unwanted species in the Ordinance to the Federal Act on Fisheries (*N. melanostomus, N. kessleri, N. fluviatilis, N. gymnotrachelus and Proterorhinus semilunaris*; (Schweizerischer Bundesrat 24.11.1993)). Further, the Strategy to combat invasive species was developed and implemented (Bundesamt für Umwelt BAFU 2016).

The project was accompanied by a variety of media communications (**Fig 2**), partly initiated by media, partly initiated by the researchers. Since policy relies on values, and values are developed early in life, we also took opportunities to participate in outreach to children. Of particular interest is a popular children’s book series with one edition featuring gobies among other invasive species (Bieri 2018).

### Learnings and observations

The output listed above suggests that the project was productive across the categories of scientific publications, career stepping stones, applied results, policy outcomes, and communication products. The process qualities of participation, reflection, and iterative adjustments were integral to the project, 1st order effects such as learning and capacity building were achieved, 3rd order effects such as influence on laws, regulations, and public discourse could be observed (Wiefek 2024), and integration of previously unrelated elements took place (Pohl 2021). Thus, we postulate that the project can be considered an example of a successful inter- and transdisciplinary project.

In Table 1, we summarize conditions which, from our perspective as natural scientists and drivers of the project, allowed the project to unfold the way it did, and allowed the researchers on the project to achieve personal academic goals while conducting meaningful, impactful research with translational value. Conditions are arranged from large-scale overarching political aspects down to the level of the individual project team member (see also Deutsch 2024 for a similar analytical framework), and combined with specific examples. Some additional observations are further explored below. Importantly, all observations and conditions were derived independently of the respective literature on id-td projects, but found to overlap substantially with previously identified facilitating conditions. Inadvertedly, our agnostic laywomanship with regard to the social science literature on id-td processes allowed for a somewhat unbiased perspective.

#### Mutual learning required time and space

An interdisciplinary project goes beyond including scientists from the different disciplines. While field-specific skills are often essential, the holders of these skills need to have the will and expertise to share their knowledge and imparting their skills on others. From this follows that resources and opportunities for knowledge transfer between the disciplines must be explicitly planned for.

#### Investments into stakeholder engagement activities ultimately paid off for the individual

Building trust and establishing dialogue beyond academic actors is resource-intense and emotionally taxing, as laid out by other sources (N’Guyen et al. 2016; Larson und Williams 2009; Moon et al. 2015; Knapp et al. 2019). Also, linking to society is not required per se for excellent research, nor is it an essential career advancing skill in the natural sciences. At times, junior researchers found it difficult to reconcile the need to invest all resources into their own career aspirations with the resource demands of stakeholder engagement. Post-hoc, however, the stakeholder-related aspects of the project built solid credibility regarding the participants’ interface science skills, and the acquired competences in managing interface research and teaching were essential during the subsequent career steps for all junior team members.

#### Diversity was a two-edged sword

The team could be considered diverse: it was female-led, with a male and a female postdoc as long-term team members, two female and one male PhD student, and several field- and lab assistants over time. Natural scientists on the project had highly distinct disciplinary backgrounds, and while this was an asset, the disciplinary perspectives on science and the scientific method also created tension and disagreements. Targeted interventions of an attentive PI were essential to instill and maintain a culture of mutual appreciation. The same constellation could have easily bred frustration and turned into a nonproductive or toxic team culture without such interventions.

#### Scientific disagreement was a source of strength

The project team was certainly not always in agreement regarding priorities or experimental design, and personal interests combined with disciplinary perspectives led to intense discussions. For example, the role of aquarists was unclear for a considerable amount of time, and not everyone considered this avenue worth pursuing. Importantly, discussions were well-moderated and led to be productive, team members were free to follow their ideas even without full buy-in from all group members, and were welcome to supply evidence for their hunches. In the case of aquarists, this led to the documentation of a substantial overlap between angler and aquarist groups, and a hitherto undiscovered risk or pathway for dispersal of non-native fish (Hirsch et al. 2021).

#### In an atmosphere of safety and playfulness, failure was a viable option

Some initial research ideas were not followed up, e.g. due to limitations of resources (e.g. systematic snorkeling for the detection of goby eggs on recreational boats, boat cleaning initiative, etc.), political will (e.g. managing ballast water in inland shipping), lack of consensus among relevant stakeholders (e.g. angler app for early invasion detection), changed research interests (e.g. PCR-based large-scale diet analyses) or difficulties implementing laboratory protocols (e.g. round goby population genetics). The decision to drop an approach or project, sometimes despite previous resource investments, was never held against individuals, and was considered a natural part of a creative process that yields more ideas than can be realized. Such a “failing is ok” approach prevented investments into “dead projects”, and ensured a focus of limited resources on the most promising ideas. Team members experienced an atmosphere of trust, transparency, and mutual reliance and commitment, also because great attention was given to the hiring of scientifically complementary but socially fitting candidates who displayed proactiveness, curiosity, and commitment to the greater cause.

#### Systemic factors were impactful

As a unit designed to produce interface content in teaching and research, the host institute was administratively located in-between research units, and faced systemic challenges such as unclear supervision rights for students. Specifically, team members were not authorized to supervise biology or environmental science master theses from the same department, which was a systemic disadvantage regarding access to motivated and capable students. Also, the small-scale non-high-throughput sequencing submissions associated with the project were back-tracked for months at the sequencing facility, which necessitated shifts in project priorities and led to “orphaned” project branches.

#### Reducing self-referentiality was crucial

Applied local studies and conservation projects with high project-specificity of results present with challenges regarding scientific exchange. Their relevance beyond the affected geographic area is often limited, few research groups conduct comparable research, and usually there are no or few pre-existing frameworks for (international) exchange, e.g. global organismic meetings or topical societies. This required the proactive creation and investment into exchange opportunities (e.g. through mini-symposia and targeted meetings with specific other research groups).

#### Personal factors and character properties played an essential role

The team was characterized, across hierarchical levels, by an openness towards unknown and novel approaches, and a willingness to leave personal comfort zones. Maintaining this “beginner’s mind” became more challenging later in the project, when the need for productive research and data analysis to reach personal career goals became more prevalent. Another prevailing characteristic of all individuals was playfulness, which ensures that activities are ‘highly-interactive’ (van Vleet und Feeney 2015).

## Discussion

This publication inspects an inter- and transdisciplinary research project in invasion biology from the perspective of the natural scientists driving it. Our aim was not to conduct a formative evaluation sensu *real-world laboratory* approaches, but to reflect on the conditions and practices that, in our view, facilitated the project’s productivity and impact on levels that we can evaluate. We use the term *facilitating conditions* deliberately to emphasize their probabilistic, non-deterministic character: they increase the likelihood that a project can generate both academic and societal value, but they do not guarantee it.

The literature on inter- and transdisciplinary research, much of it from the social sciences, has already identified a wide array of “success factors” and “facilitating conditions” (e.g. Pohl 2021, Wiefek 2024, Deutsch 2025). What our contribution adds is an inside view from natural scientists embedded in such a project. This perspective highlights dimensions that may be overlooked when projects are assessed from the outside. This includes the tension between career needs and project demands, especially for early-career researchers who must balance stakeholder engagement with disciplinary output and may not be able to voice or discuss these struggles openly; the role of disciplinary disagreement as creative driver, not just risk to be managed; the lived experience of systemic constraints (e.g. thesis supervision rights, sequencing bottlenecks) that rarely surface in higher-level analyses but profoundly shape feasibility; and unpleasant realizations that team diversity may be necessary in areas where one may wish it was not: gender played a role when establishing reports with male-dominated stakeholder groups such as angler associations. These are not abstract “factors” but grounded, sometimes uncomfortable realities. They show why natural science voices are essential in the broader discourse on inter- and transdisciplinary research.

Our case features facilitating conditions that span three domains: external settings (political culture valuing evidence, approachable administrations, pre-existing societal networks, and diverse funding landscapes), internal practices (deliberate management of diversity, tolerance of failure, flexible resource allocation, proactive networking, and career-sensitive authorship strategies) and personal dispositions (curiosity, openness, and the willingness to step outside disciplinary and personal comfort zones). The interaction of these domains created “perfect storms of opportunity,” where momentum emerged not from a single factor but from their convergence. This echoes frameworks in the social sciences (Deutsch 2025) but grounds them in lived experience.

It goes without saying that the facilitating conditions we describe are not universally transferable. For example, in contrast to the aquatic setting of the described project, terrestrial stakeholder-, management- and policy landscapes focus less on habitats and more on species (groups), and are thus more fragmented (Heiderich et al. 2024), which complicates truly integrative (Pohl 2021) approaches. Also, own exchanges with neighboring countries’ representatives suggest fundamental differences in terms of challenges and opportunities when decision structures are highly centralized. Also, some of the conditions listed must be considered priors with limited or no opportunities for active shaping by a research team. For example, it is beyond the individual research group to establish a political environment that provides trust and resources for investigation. Nevertheless, being explicit about such conditions can help teams anticipate opportunities, mitigate constraints, make intentional strategic choices and capitalize on strongholds, and thus may increase overall chances for a productive and positively perceived project.

For social science research around id-td research, we want to conclude with the suggestion to (also, on occasion) write for the audience who most often instigates and drives id-td projects: natural scientists. Engaging natural scientists not only as objects of research but also as co-authors and peer commentators may ensure that lessons learned are both analytically rigorous and practically useful. An excellent example for resources aimed at the natural science community is provided by the td-net of the swiss academies of sciences (scnat). Additionally, efforts could be geared towards sharing the immense expertise of social sciences in the context of natural scienc graduate schools or masters programs.

For natural scientists, our key recommendation is simple: treat reporting on facilitating conditions as part of the project itself. Collecting indicators across outputs, policy links, and engagement processes provides evidence of societal impact, which is increasingly demanded under frameworks such as DORA, and strengthens the case for inter- and transdisciplinary research as a credible academic pathway. Public expectations for demonstrating societal impact of research are growing (Filchenko et al. 2024), and “soft” indicators are increasingly integrated into institutional evaluation processes. Systematic reporting - from grants to media outputs - is becoming an essential part of justification for research units at the interface of science and society. We would also like to encourage natural scientists in id-td research projects to take the time for reflection, and provide the community with examples and observations similar to this account.

## Acknowledgements

We would like to express our gratitude towards several colleagues for critically reading and commenting on the manuscript draft, to all funders and supporters, to essential reviewer’s comments that much improved the manuscript, and to all stakeholders who participated in the project.

## Footnotes

1 applicable exchange rate: 1 CHF ~ 1 EUR ~ 1.2 USD

